# Reduction of a-synuclein aggregates by PIKfyve inhibition via TFEB-mediated lysosomal biogenesis in a Parkinson disease model

**DOI:** 10.1101/2023.10.23.563557

**Authors:** Sara Lucas-Del-Pozo, Giuseppe Uras, Federico Fierli, Veronica Lentini, Sofia Koletsi, Kai-Yin Chau, Derralynn Hughes, Anthony HV Schapira

## Abstract

Parkinson disease is a neurodegenerative disorder characterised by impairment of motor function, and is associated with a progressive accumulation of insoluble aggregates of misfolded alpha-synuclein. In the present study, we exploited the SH-SY5Y cell model overexpressing a pro-aggregation form of alpha-synuclein to investigate the efficacy of PIKfyve-mediated lysosomal biogenesis, through TFEB, as potential target for Parkinson therapy.

To investigate this, we exploited high-content imaging along with enzymatic assays to follow the progression of lysosomal biogenesis, lysosomal function and alpha-synuclein accumulation. The cellular model exploited in this study recapitulated important elements of the biochemical phenotype observed in Parkinson patient-derived neurons, including synuclein aggregates and impaired glucocerebrosidase (GCase) function. PIKfyve inhibition by YM201636 resulted in a lysosomal-dependant reduction of alpha-synuclein aggregates as early as 24 hours post-treatment. The mechanism of action of YM201636 was shown to be TFEB-mediated, with an increase in TFEB in the nuclei which subsequently resulted in increased lysosomal markers LAMP1 and GCase. PIKfyve inhibtion efficacy was also tested in differentiated SH-SY5Y cells, exhibiting a neuron-like morphology. In these cells, YM201636 also significantly reduced alpha-synuclein aggregates and increased TFEB nuclear presence.

These findings suggest that PIKfyve inhibition could be exploited as therapeutic target for Parkinson disease.

## Introduction

Parkinson disease (PD) is the second most common neurodegenerative condition after Alzheimer disease (AD) ^1^. The main risk factor for developing PD is age, affecting about 3% of the population from 65 years ^2^, with more than 6 million individuals affected currently worldwide. Histopathologically, PD is defined by the presence of cytoplasmic protein inclusions called Lewy bodies, whose main component is the protein alpha-synuclein (a-synuclein) ^3,4^. Furthermore, in these neuronal a-synuclein enriched inclusions (Lewy bodies and Lewy neurites), both misfolded and hyperphosphorylated a-synuclein at serine129 (pS129)^5^ as well as a lipid-enriched fraction (including vesicular structures and dysmorphic organelles) are found^3^. Accumulation and aggregation of misfolded a-synuclein is associated with a progressive loss of dopaminergic neurons in the substantia nigra pars compacta (SNpc), and a reduction of dopamine levels in the striatum.

Although the aetiology of PD is not fully known, both genetic and environmental factors influence PD susceptibility. Variants in the *GBA1* gene which encodes the lysosomal enzyme glucocerebrosidase (GCase), represent the most important genetic risk factor for PD ^6,7^. GCase is a membrane-associated lysosomal hydrolase that catalyses the cleavage of glucosylceramide (GlcCer) and glucosylsphingosine (GlcSph) into glucose and ceramide, and glucose and sphingosine, respectively. In the biallelic or heterozygous state, *GBA1* genetic variants increase the risk for PD ^6^. Up to 1% of individuals in the general population are non-manifesting heterozygous *GBA1* carriers and they have up to a 20-fold increased risk to develop PD as compared to non-carriers ^8–11^.

Low levels of GCase activity have been reported in post-mortem brain from people with sporadic PD without *GBA1* genetic variants^12,13^ and several studies showed that impaired GCase activity results in increased a-synuclein levels in cell and animal models^14,15^ as well as in human induced pluripotent stem cells (hiPSC) ^16,17^, linking lysosomal function to PD pathogenesis ^18,19^.

The autophagic and endolysosomal pathways are crucial for a-synuclein elimination^20^. A-synuclein accumulation causes lysosomal dysfunction in PD-patient-derived iPSC-midbrain neurons^21^ and reduces lysosomal degradation in human midbrain dopamine models of synucleinopathies through disrupting GCase trafficking from the endoplasmatic reticulum (ER) to the lysosomes ^22,23^. It has been reported that a-synuclein accumulation at the ER, causing ER fragmentation, compromising folding capacity and aggregation of lysosomal hydrolases in the ER ^24^. Consequently, enhancing lysosomal and potentially GCase activity as a treatment strategy to slow progression of a-synuclein pathology has gained attention during the last decade.

Transcription Factor EB (TFEB) is a helix-loop-helix (HLH) leucine-zipper transcription factor capable of recognising sequences of the Coordinated Lysosomal Expression and Regulation (CLEAR) gene network, which coordinates lysosomal biogenesis and function^25^. TFEB positively regulates the expression of lysosomal biogenesis-related genes, resulting in an increased number of lysosomes ^26^ and higher levels of lysosomal enzymes, eventually enhancing lysosomal catabolic activity ^27^. TFEB also upregulates autophagy and lysosomal exocytosis, further promoting autophagosome formation, autophagosome-lysosome fusion, and degradation of autophagy substrates, enhancing the clearance of intracellular misfolded proteins ^28–31^. TFEB is constitutively found in its phosphorylated state in the cytoplasm attached to the lysosome. Upon de-phosphorylation of multiple serine residues (mainly Ser142, Ser211), TFEB re-locates to the nucleus, thereby exerting its transcriptional activity ^32^. Mechanistic target of rapamycin complex 1 (mTORC1) is the most relevant kinase known to phosphorylate TFEB, playing a crucial role in the regulation of TFEB subcellular localisation by phosphorylating Ser211, promoting TFEB retention in the cytoplasm. Different signals are capable of positively regulating TFEB translocation into the nucleus, such as starvation and cell division ^27,32^. Increasing lysosomal biogenesis and catabolism via TFEB upregulation is therefore a potential pathway to further clear misfolded proteins, already studied in neurodegenerative diseases, such as PD and Alzheimer’s disease (AD) ^33^ ^34^.

A regulator of TFEB translocation is lipid kinase PIKfyve, which was first identified over two decades ago as a lipid kinase that phosphorylates phosphatidylinositol-3-phosphate(PI(3)P), producing PI(3,5)P2. It is named after its function and domain structure - phosphoinositide kinase for five position containing a FYVE finger-and contributes to regulating membrane homeostasis, endosomal trafficking and autophagy ^35^. In order to carry out its kinase activity, PIKfyve must form a complex that includes the scaffold protein Vac14 and the lipid phosphatase Fig 4 ^36^.

PIKfyve plays a crucial role in the maintenance of ion homeostasis in lysosomes via its product PI(3,5)P2, which binds to key lysosomal ion channels including the mucolipin TRP channel (TRPML1) ^37^ and also contributes to maintain the lysosomal acidic pH ^38^. Membrane dynamics in the endolysosomal system are also tightly regulated by PIKfyve activity since it is essential to drive lysosome reformation from endolysosomes ^39^. PIKfyve is necessary for the generation of early endosomes ^40^ and it also plays a role in endocytic recycling to the plasma membrane ^41^.

In mammalian cells, PIKfyve also plays roles in the regulation of transcription, as PIKfyve is a key regulator of TFEB. PIKfyve acts by positively regulating mTORC1-dependent phosphorylation of Ser-211 on TFEB ^42^. Upon PIKfyve inhibition, protein phosphatase 2A dephosphorylates TFEB Ser-211, which results in nuclear translocation of TFEB^43^.

The initial inhibitor identified for PIKfyve was YM201636, a pyrimidine-based kinase inhibitor with good potency against PIKfyve *in vitro* and selectivity over several lipid kinase family members ^44^ ^35^. The efficacy of PIKfyve inhibition as a potential therapeutic intervention for neurodegenerative conditions such as Parkinson’s has been highlighted in genetic screening, in which PIKfyve inhibition prevented an increase of a-synuclein aggregation^45^. However, such effects are unexpected given research indicating that decreased PIKfyve activity in mouse models produce a spongiform brain pathology ^46^. Mutations in the PIKfyve complex are associated with neurodegenerative diseases such as Charcot-Marie-Tooth syndrome ^47–49^(CMT4J) and amyotrophic lateral sclerosis (ALS) ^50,51^, among others. However, the human disease recognised to be associated with mutations in the PIKfyve gene is Fleck corneal dystrophy, a rare autosomal dominant disease of the corneal stroma characterized by intracytoplasmic accumulation of glycosaminoglycans and complex lipids in keratocytes ^52,53^. The fact that this hereditary corneal dystrophy is not associated with neurodegeneration indicates that partial reduction of PIKfyve activity is viable and well tolerated.

To investigate whether the activation of TFEB signalling through PIKfyve inhibition can prevent the formation of a-synuclein aggregates, we exploited a cellular model overexpressing a mutant form of the *SNCA* gene which leads to a rapid aggregation of a-synuclein protein following its interaction with lipid-membrane organelles ^54^. In this mutant model, the familiar PD mutation E46K ^55^ was “amplified” by introducing two analogous extra-lysine mutations into the nearby KTKEGV repetitive motive in positions E35K, E46K, and E61K ^56^. The presence of such mutations leads to a rapid aggregation of a-synuclein, detectable after 24-48 hours in culture ^54^.

Here we report that treatment with the PIKfyve inhibitor YM201636 results in a significant reduction of a-synuclein aggregates in neuroblastoma cells overexpressing the 3K mutation. The effect is lysosomal-dependent, since the amelioration observed was prevented when bafilomycin, which impairs lysosomal function, was administered together with this compound ^57,58^. In the study described here, we also show how YM201636 treatment reduces a-synuclein aggregates following differentiation of the neuroblastoma cells into neuron-like cells, with an overlapping mechanism of action through TFEB activation. We propose TFEB activation, and subsequent increase in lysosomal levels, as a potential target pathway for the clearance of a-synuclein aggregates.

## Methods

### Cell line and compounds

SH-SY5Y cells were used as parental control cells (#91396-88-2, Atcc,UK). SH-SY5Y cells constitutively expressing the 3K-SNCA gene were a kind gift from Dr. Ulf Dettmer laboratory (Harvard, USA). Compounds YM201636 (#13576, Cayman Chemical, Ann Arbour, US) and ambroxol hydrochloride (#A9797, Merck, London, UK) were diluted in DMSO to obtain a 10mM stock solution. Subsequent dilutions were all made in sterile PBS and stored at −80C until usage.

### Cell culture

SH-SY5Y cells and 3K-SNCA overexpressing cells, were maintained in 1:1 DMEM/F12 media with Glutamax (Thermo, Waltham US) supplemented with 10% fetal bovine serum (FBS) (Thermo, Waltham US) and 1X Penicillin/Streptomycin (Thermo, Waltham US). Cells were passaged appropriately to maintain a confluency between 50-70%.

### Neuronal differentiation

Wild-type and 3K-SNCA overexpressing SH-SY5Y were differentiated into neurons using retinoic acid-based protocol. 70% confluent cells were detached using trypsin and Versene® and resuspended in Neurobasal media (Thermo, Waltham US) supplemented with 30μM retinoic acid (Thermo, Waltham US), 1X vitamin B-27 (Thermo, Waltham US), 1X Glutamax (Thermo, Waltham), and 5ng/ml brain-derived neurotrophic factor (BDNF) (R&D Systems, Abingdon UK). Differentiation medium was replaced every 48 hours, with a neuronal morphology appearing after 4 days of culture. Differentiated neurons were used for downstream analysis after 10 days in culture. Cells were seeded in Geltrex® Matrix solution diluted in DMEM/F-12 medium (1:100) coated plates at 1.5 × 10^5^ cells/ml.

### Compound treatments

Non-differentiated cells were grown up to 70% confluency on standard cell culture media. Subsequently, cells were detached using trypsin, counted, and seeded in a new 96-well plate. 3*10^4^ cells were seeded into each well, avoiding placing them into borders, rows and columns to prevent plate-edge effects. 24 hours post seeding, the medium was discarded and replaced with 100μl of medium containing the appropriate compound or vehicle. Cells were then left to incubate for 24 hours and subsequently used for downstream analysis.

Differentiated cells in culture for 9 days were treated with 100μl of medium containing the appropriate compound, or vehicle, for 24 hours and subsequently used for downstream analysis.

### Cell staining

Following compound treatment for 24 hours, cell culture medium was removed and cells fixed with 100μl of 4% para-formaldehyde (PFA) (Merck, London UK) for 20 minutes at room temperature. Subsequently, PFA was removed and 100μl of ice-cold methanol was added to each well for 20 minutes. Methanol was then discarded and each well washed 3 times with 1X PBS (Thermo, Waltham US). Cells were then blocked with 100μl 5% normal goat serum (NGS) (Thermo, Waltham US) for 1 hour at room temperature. Primary antibodies were then added according to the experiment: monoclonal rabbit anti-synuclein 1:750 (#ab138501, Abcam, Cambridge UK), monoclonal mouse anti-GBA 1:1000 (Anti-GBA mAb (2E2), #AP1140-100UG), monoclonal rabbit anti-TFEB 1:500 (#4240, Cell Signalling, Danvers US), monoclonal mouse anti-LAMP1 1:1000 (#AB_398356, BD Biosciences, US). Upon addition of primary antibody, cells were left to incubate overnight at +4C. Cells were then washed 3 times with PBS and incubated for 2 hours at room temperature with secondary antibody according to the experiment: AlexaFluor 488 1:2000, AlexaFluor 564 1:2000, AlexaFluor 647 1:2000. Subsequently cells were washed 3 times with 1X PBS, and incubated for 10 minutes with 100μl of DAPI solution (5μg/ml). Each well was then washed 3 times with 1X PBS and filled with 200μl of PBS. Plates were then stored at +4C avoiding light until analysis.

### High content imaging

Cell imaging was performed using Opera Phoenix Plus HCS (PerkinElmer, Waltham US) using confocal mode. Each channel was acquired separately to avoid overlap signal. DAPI was acquired with laser power at 30% with exposure set at 100ms. 488 channel was acquired with laser power at 70% with exposure at 100ms. 564 channel was acquired with laser power at 70% with exposure at 100ms. 647 channel was acquired with a laser power at 80% with exposure at 100ms. For each image a z-stack of 2μm was acquired.

Image analysis acquisition was done using Harmony^TM^ Phenix software and image analysis was performed using Columbus™ Image Data Storage and Analysis system (PerkinElmer, Waltham US). Each z-stack was first converted into a maximum projection image before proceeding with further analysis. Nuclei were first identified using built-in method B in Harmony. Cytoplasm was then identified using Nuclei as reference point, and built-in method A in Harmony to identify the extent of the area. To identify a-synuclein aggregates, a filter mask to the 488 channel in the cytoplasm area was applied, with a signal intensity range between 5500 and 20000, getting as a result the number of aggregates per field. Data were then normalised by the number of nuclei, therefore quantifying the number of a-synuclein aggregates for each single cell. To quantify TFEB partitioning between cytoplasm and nuclei, a filter mask to the 568 channel was applied in both areas, with a signal intensity range between 150 and 20000. The mean fluorescent intensity of TFEB in the nuclei area in each well was obtained through the Calculate Intensity Properties feature built-in method in Harmony, and then it was normalised by number of nuclei, obtaining the mean nuclei TFEB intensity per cell in each field. GCase intensity per cytoplasm was quantified using a filter mask in cytoplasm on channel 568, with a signal range between 600 and 20000. The mean fluorescent intensity of GCase in the cytoplasmatic area in each well was obtained through the Calculate Intensity Properties feature built-in method in Harmony, and then it was normalised by number of nuclei, obtaining the mean cytoplasmatic GCase intensity per cell in each field.

### HTRF HUMAN - alpha synuclein aggregation assay

The quantification of synuclein aggregates was performed using HTRF alpha-synuclein aggregation kit according to manufacturer instruction (#6FASYPEG & #6FASYPEH, Cisbio, London UK). Cell samples were first diluted in water. Subsequently, 5μl of Lysis buffer, along with 5μl of both Anti-h-s-Synuclein-d2 antibody and Anti-h-a-Synuclein Tb-Cryptate antibody, were added to the sample and left to incubate for 20 hours at room temperature. Fluorescence emission was then read at both 665nm and 620nm. A ratio between 665nm/620nm was then applied to quantify the aggregated synuclein.

### GCase activity assay

A total of 3*10^4^ cells were seeded in each well of 96-well plate and left to incubate for 24 hours. For differentiated cells, day 9 cultured cells were used. Subsequently, the medium was replaced with compound containing medium and incubated for 24 hours. Cell medium was then removed and each well washed 2 times with 1X PBS at room temperature. Cells were then lysed using 50μl cold lysis medium (50mM citric acid, 0.15M K_2_HPO_4,_ 10mM Sodium Taurocholate, 0.01% Tween-20, 20% Triton X, and 1X protease inhibitor cocktail). Subsequently, 50μl of buffer (50mM citric acid, 0.15M K_2_HPO_4,_ 10mM Sodium Taurocholate, 0.01% Tween-20, final pH5.9) supplemented with 1mM 4-MUG (#M3633, Merck, London UK) were added to each well and samples left to incubate overnight at room temperature protected from light. Reaction was then stopped by adding 100μl of stop solution (0.5M NaOH, 0.5M glycine) to each well, and then left to incubate for 1 hour at room temperature. Plates were then analysed using Cytation^TM^ plate reader using and endpoint protocol with wavelength ex/em set up at 360/460nm. Obtained data was then normalised to control and expresses as a fraction.

### Data analysis

Statistical analysis was performed using GraphPad Prism^TM^ 9.1 software. Each experiment was carried out in triplicate or duplicate for each condition. Baseline results of quantitative variables are expressed as mean ±standard error of the mean or ±standard deviation for each categorical group. In order to make comparisons, the Shapiro-Wilk W test was first used to determine whether the collected data followed a normal distribution. In non-normally distributed data, multiple categorical groups were compared by ANOVA with Bonferroni correction (parametric test) or Kruskal-Wallis with Dunn’s post-hoc analysis (non-parametric) for quantitative dependent variables. Results were deemed statistically significant if their p-value was less than 0.05.

## Results

### Overexpression of 3K-SNCA mutant gene leads to formation of a-synuclein aggregates and lysosomal dysfunction

To understand whether overexpression of *3K-SNCA* leads to a rapid formation of visible aggregates of a-synuclein in the cells, we compared the amount of a-synuclein aggregates per cell in *3K-SNCA* mutant SH-SY5Y cells to wild-type SH-SY5Y. High content imaging analysis showed not only an increased mean a-synuclein signal intensity in the cell body of *3K-SNCA* cells, likely due to the constitutive overexpression, but also a statistically significant increase in the number of a-synuclein aggregates per cell as early as 24 hours post-seeding (Fig.1 a,b) when compared to wild-type cells. The quantification performed revealed that *3K-SNCA* cells develop at least one a-synuclein aggregate in their cytoplasm, as opposed to the control group, which shows less than 0.01 aggregates per cytoplasm. These results recapitulate the increase in a-synuclein inclusion formation in *3K-SNCA* cells previously described in the literature ^54^. To further confirm this, we performed a FRET-HTRF assay to quantify the levels of aggregates. Our results, show an increase of about 10-fold in the *3K-SNCA* overexpressing cells (Fig. S1), in line with the findings obtained using confocal analysis.

**Figure 1.**
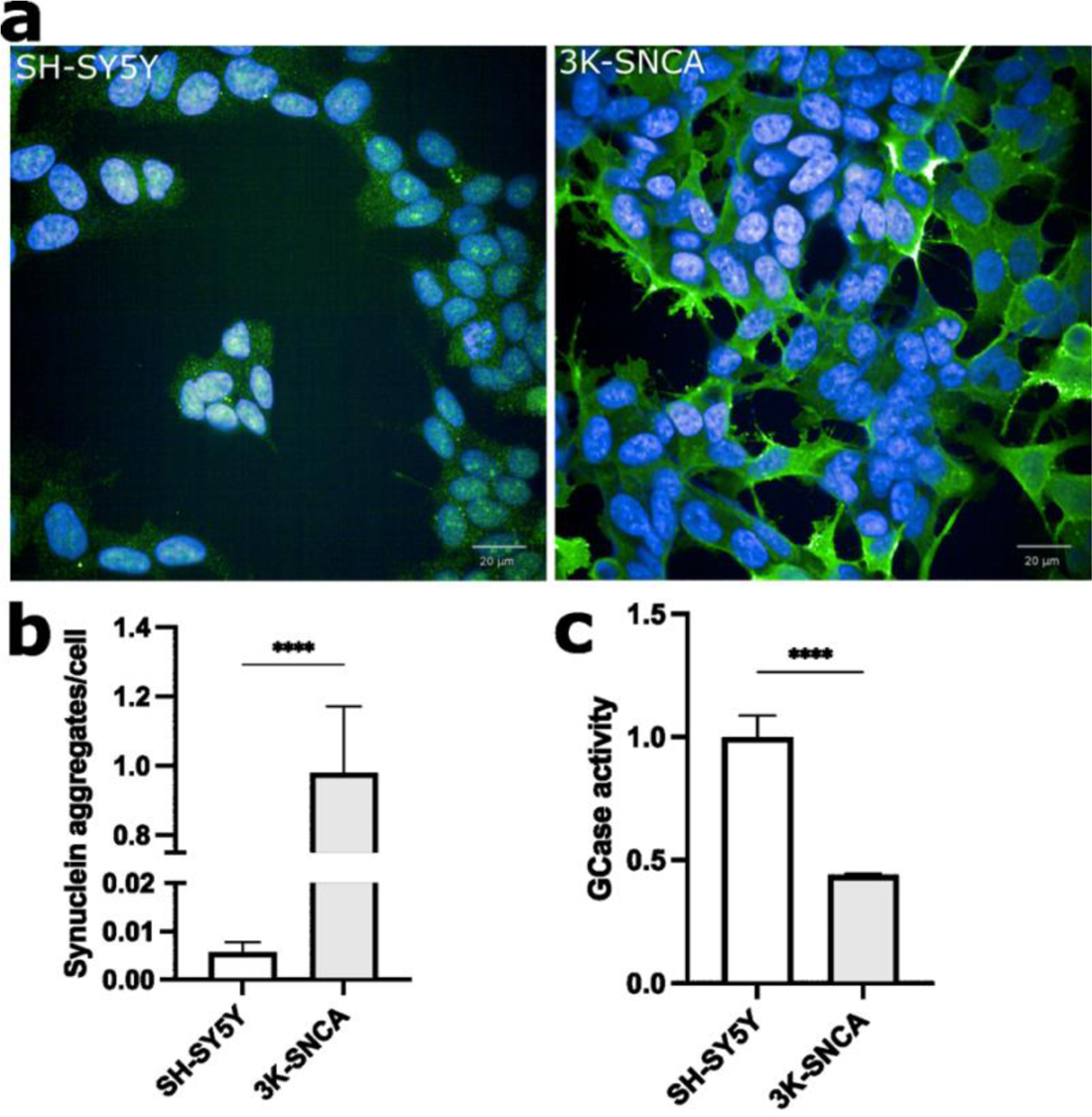
Characterization of SH-SY5Y cells overexpressing *3K-SNCA* mutant gene. **a)** Representative confocal images showing a-synuclein aggregates spots in *3K-SNCA* expressing cells compared to wild type SH-SY5Y with weak α-synuclein signal. **b)** Quantification of a-synuclein spots in *3K-SNCA* cells and wild type SH-SY5Y. **c)** Quantification of total GCase activity in *3K-SNCA* and wild type SH-SY5Y. Data are presented as mean±SEM of 3 independent experiment. ***=*P<*0.001; *\*\*\*\**=*P*<0.0001.

Following the identification of a significant increase in a-synuclein aggregate number in *3K-SNCA* expressing cells, we investigated whether the presence of such aggregates would lead to lysosomal dysfunction. Specifically, due to the reciprocal relationship previously described between a-synuclein and GCase^59^, we focused on this particular hydrolase. It has been documented that increased a-synuclein levels lead to a reduction in wild-type GCase activity and protein expression ^60^.

Therefore, we quantified the total activity of GCase enzyme in cell lysates using fluor-labelled substrate 4-MUG to study whether wild-type GCase activity was impacted in our cell model. Endpoint analysis of total GCase activity revealed an approximate 50% reduction in enzyme activity in *3K-SNCA* expressing cells when compared to the wild-type group (Fig 1c). This result confirms that overexpression of mutant a-synuclein and formation of aggregates compromises wild-type GCase activity, recapitulating previous results in a-synuclein overexpressing models in which enhanced a-synuclein aggregation and accumulation result in selective reduction in GCase lysosomal hydrolase activity in synucleinopathies ^24,60^.

### PIKfyve inhibitor reduce synuclein aggregates via lysosomal-dependant pathway

We tested the effects of the PIKfyve inhibitor YM201636 on a-synuclein accumulation in the 3K-SNCA cells. Initially, to identify the optimal drug dosage, we performed a pilot experiment treating the *3K-SNCA* overexpressing cells with serial concentration of the PIKfyve inhibitor YM201636 for 24hours, using as readout the total number of a-synuclein aggregates per cell. Our results indicated that YM201636 treatment, although not reaching statistical significance, displayed a trend towards a decrease of a-synuclein aggregates at 1uM (Fig. S2). Subsequently, *3K-SNCA* cells were treated with YM201636 at the specified concentration and a vehicle control for 24 hours. Ambroxol, an oral mucolytic drug that has been used since late 1970s, acts as a pH-dependent, mixed-type inhibitor of GCase ^61^.

Additionally, ambroxol is allegedly capable of modulating TFEB ^62,63^.Consequently, ambroxol control treatment was administered at 30uM as previously reported ^19^ resulting in a 20% decrease. PIKfyve inhibition by YM201636 produced a more effective reduction, with the number of a-synuclein aggregates being reduced by 50% (Fig.2; a,b). This indicates that a-synuclein accumulation is partially reversible both upon TFEB stimulation.

**Figure 2.**
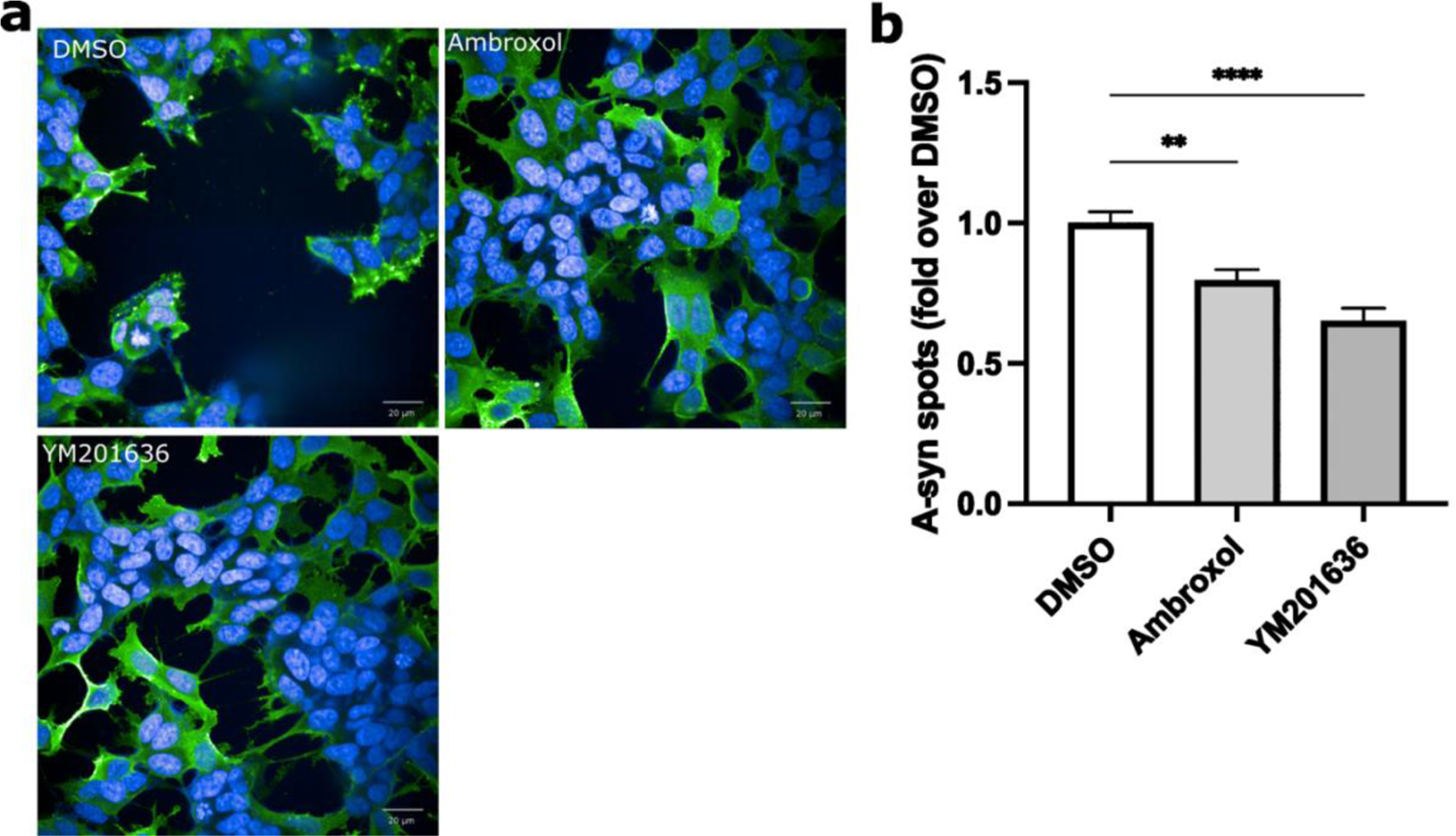
Compounds-driven reduction of synuclein aggregates in 3K-SNCA cells after 24 hours treatment. **a)** Representative confocal images showing reduction of synuclein aggregates in 3K-SNCA treated cells compared to vehicle control. **b)** Quantification of synuclein spots in 3K-SNCA cells following treatment. Data represented as fold-change over vehicle control. Data is presented as mean±SEM of 3 independent experiment. **=*P*<0.01; *\*\*\*\**=*P*<0.0001.

Finally, since YM201636 has been described to induce the nuclear translocation of TFEB ^43^, we investigated whether: (i) TFEB stimulation effects are exerted, at least to some extent, through the lysosomal system, and (ii) a constitutively healthy lysosomal function is indispensable to induce a reduction of a-synuclein aggregates under TFEB stimulation. We treated *3K-SNCA* overexpressing cells with bafilomycin-A1, a compound that de-acidifies the lysosome in an indirect way by inhibiting the vacuolar-type ATPase in the lysosomal membrane and impairing lysosomal function ^57^. Confocal image analysis revealed that no significant reduction of a-synuclein aggregates was achieved by YM201636 or ambroxol when co-administrated with bafilomycin-A1 (Fig.3; a,b). These findings indicate that the reduction in a-synuclein induced by YM201636 is mediated through the lysosome.

**Figure 3.**
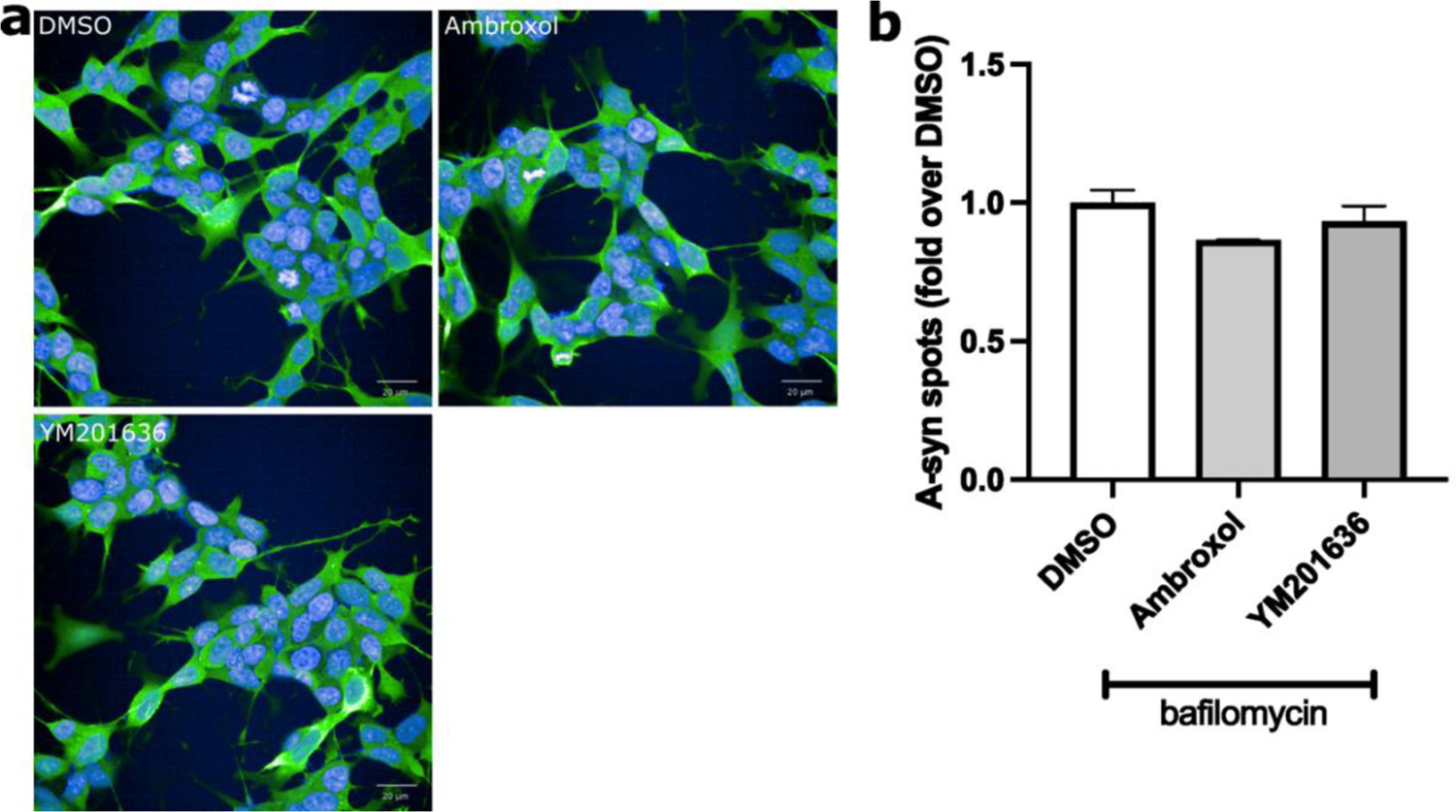
Co-treatment of bafilomycin-A1 and compounds on *3K-SNCA* expressing cells. **a)** Representative confocal images showing inhibition of reduction of a-synuclein aggregates for ambroxol and YM201636 following impairment of lysosomal function. **b)** Quantification of synuclein spots in *3K-SNCA* cells following co-treatment with 10nM bafilomycin for 24 hours. Data is presented as mean±SEM of 3 independent experiment.

### PIKfyve inhibition treatment increase TFEB translocation into the nuclei and subsequent lysosomal biogenesis

Our data suggest a lysosomal-dependant reduction of a-synuclein aggregates following treatment with PIKfyve inhibitor YM201636 and control drug ambroxol. Consequently, we hypothesised that the rescue of the a-synuclein aggregation and accumulation is driven by an upregulation of TFEB-dependant pathways. We first investigated whether an increase in TFEB levels was detectable following compound treatment at different time points. We found a trend towards increasing levels of TFEB signal intensity in the nuclei after 4 and 7 hours of treatment, ultimately reaching a statistically significant difference in TFEB nuclear levels when comparing with vehicle control after 24 hours of treatment for all the compounds (Fig. 4, a-d). Subsequently, to test whether the observed increased TFEB nuclear levels resulted in an increase in lysosomal density in the cell cytoplasm, we quantified GCase protein, which is located in the lysosomal lumen ^64^, using a high content imaging approach. Our analysis identified a significant GCase increment in *3K-SNCA* cells treated with ambroxol after 24 hours of treatment, mirroring the results showed on TFEB translocation. YM201636 displayed a trend towards an increase in GCase protein levels, in line with the limited increment observed on TFEB analysis (Fig. 5, a-b).

**Figure 4.**
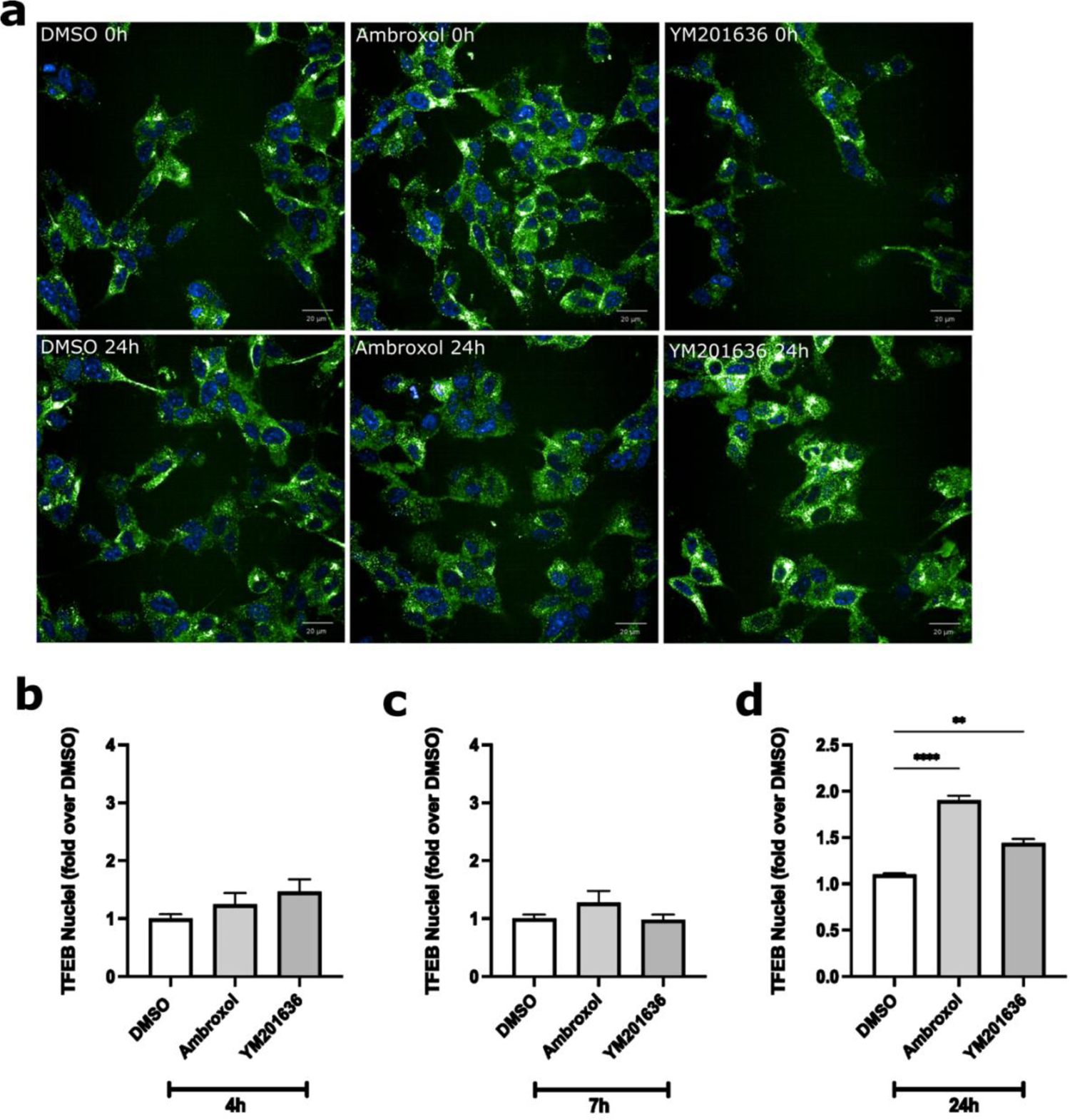
TFEB increased translocation into the nuclei following compound treatment. **a)** Representative confocal images showing increased TFEB into the nuclei after 24 hours treatment. **b)** Quantification of TFEB nuclei signal intensity after 4 hours treatment **c)** Quantification of TFEB nuclei signal intensity after 7 hours treatment **d)** Quantification of TFEB nuclei signal intensity after 24 hours treatment. Data is presented as mean±SEM of 3 independent experiment. **=*P*<0.01; *\*\*\*\**=*P*<0.0001.

**Figure 5.**
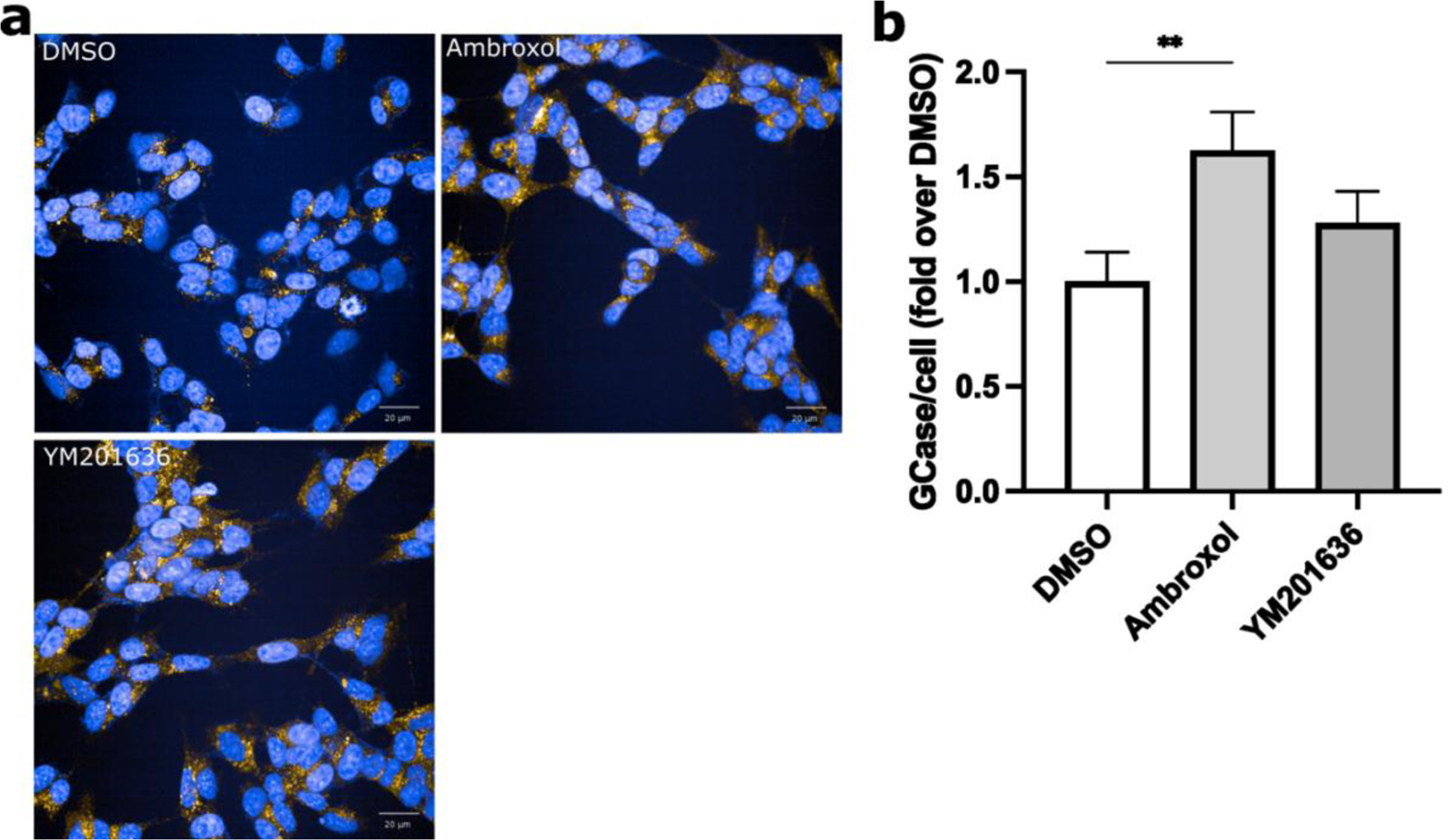
Ambroxol treatment increases GCase intensity per cell when compared to vehicle control. **a)** Representative images showing increased GCase levels after compounds exposure. **b)** Quantification of GCase after 24 hours treatment. Data is presented as mean±SEM of 3 independent experiment. *=*P*<0.05.

We, then, questioned whether this enhanced TFEB nuclear translocation and increment in GCase levels after 24 hours of treatment would increase GCase activity, potentially reversing the increased a-synuclein aggregation. Our analysis did not reveal any significant increase in whole GCase activity at this stage in any group (Fig. S3). Thus, to confirm that ambroxol and YM201636 treatments modulate not only GCase protein content, but the overall lysosomal biogenesis, we quantified the presence of the Lysosomal Associated Membrane Protein 1 (LAMP1). This lysosomal external membrane protein is specifically located in mature lysosomes and not found in other endosomal compartments ^65^. Our analysis shows how PIKfyve inhibition significantly increases LAMP1 punctate after 24 hours, confirming the trend observed on GCase content. In contrast, ambroxol treatment did not result in an overall increase in LAMP1, with effects limited to GCase levels (Fig. 6, a-b). Together, these data suggest that YM201636 reduction of a-synuclein is achieved by promoting TFEB translocation into the nucleus, ultimately increasing the pool of lysosomes. However, this is not associated with a relative increment of global GCase activity.

**Figure 6.**
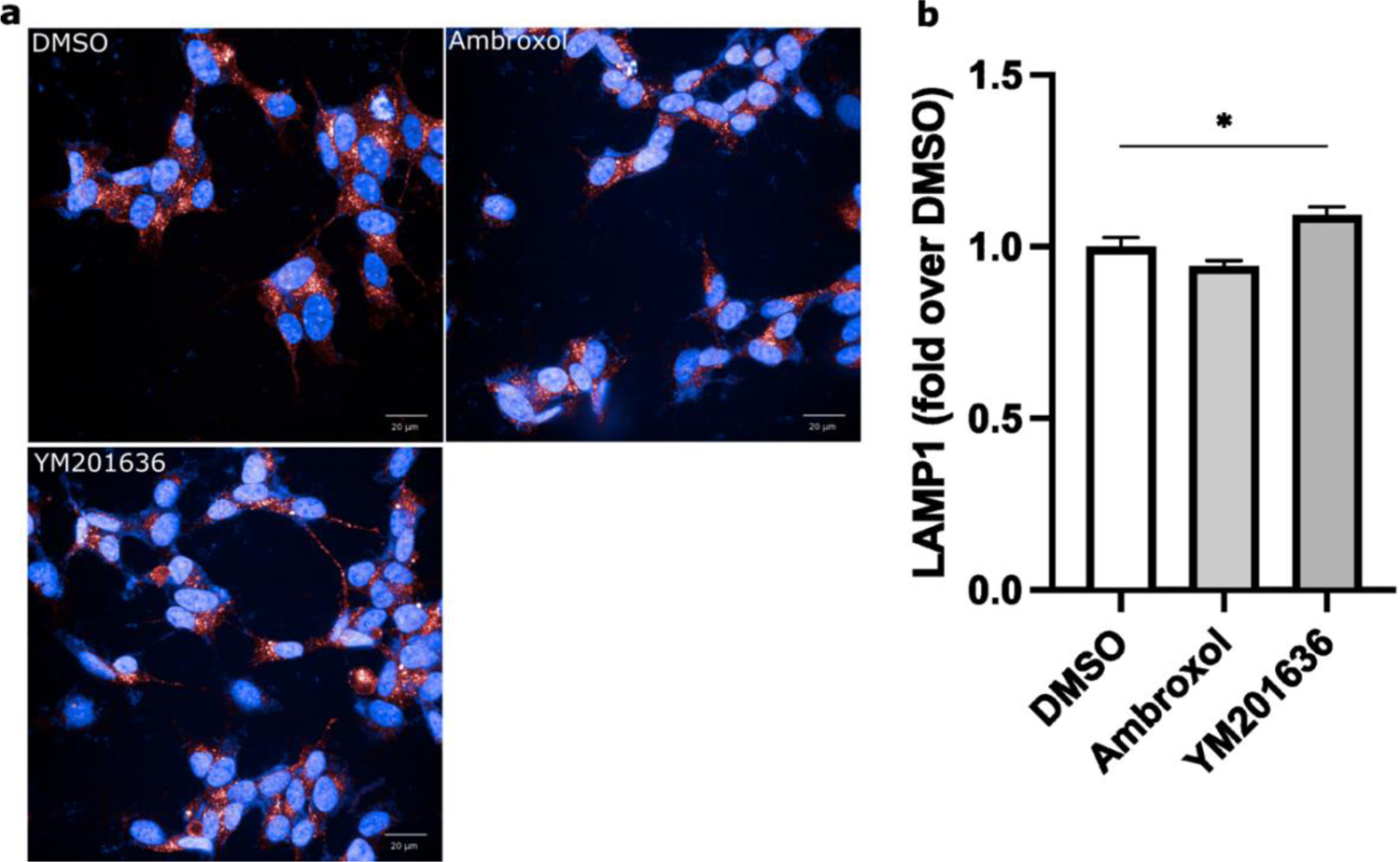
YM201636 treatment increase LAMP1 presence per cell when compared to vehicle control. **a)** Representative images showing increased LAMP1 levels after compounds exposure. **b)** Quantification of LAMP1 after 24 hours treatment. Data is presented as mean±SEM of 3 independent experiment. *=*P*<0.05.

### Inhibition of PIKfyve reduces a-synuclein burden via TFEB increase in differentiated 3K-SNCA overexpressing cells

To investigate whether our findings would be recapitulated in *3K-SNCA* overexpressing cells without active division, we obtained derivative cells with a neuronal phenotype by using a RA and BDNF induced neuronal differentiation protocol ^18^. This procedure leads to a homogeneous neuronal population with expression of neuronal markers and decreased proliferation ^66^. After 10 days in culture using media supplemented with RA and BDNF, SH-SY5Y overexpressing *3K-SNCA* cells presented a neuronal morphology (Fig. S4), still exhibiting the presence of a-synuclein aggregates (Fig. 7, a).

**Figure 7.**
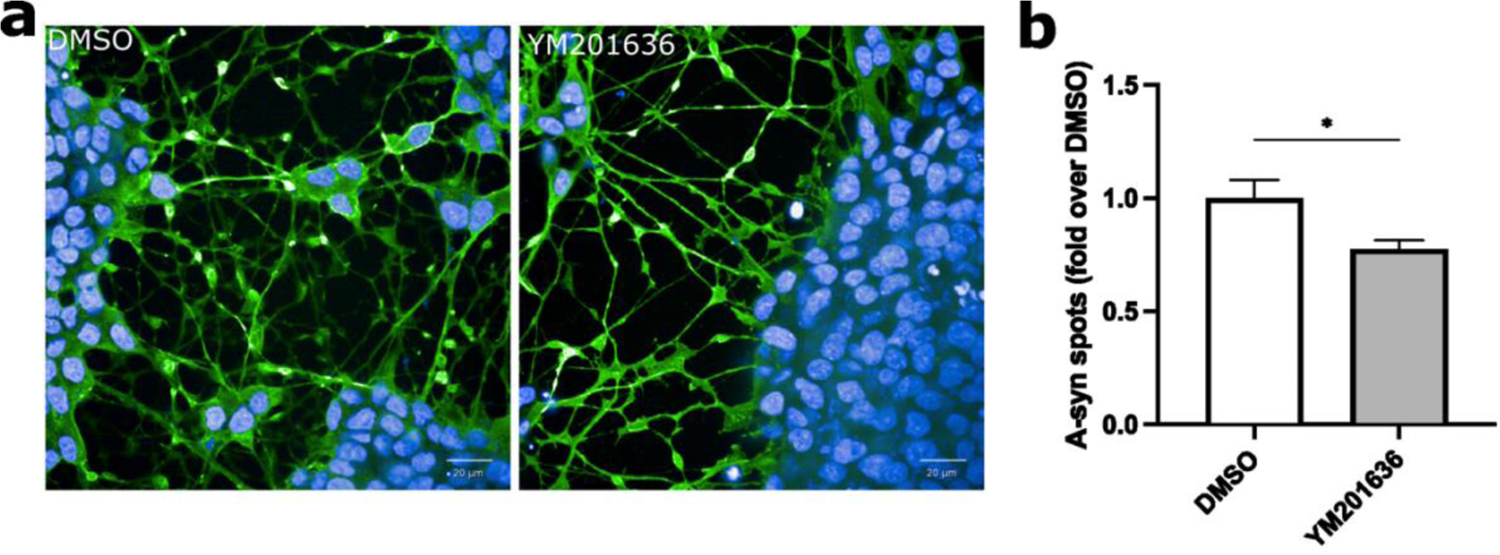
Compounds-driven reduction of synuclein aggregates neuron-like cells overexpressing *3K-SNCA* gene. **a)** Representative confocal images showing reduction of synuclein aggregates in differentiated *3K-SNCA* treated cells compared to vehicle control. **b)** Quantification of synuclein spots in *3K-SNCA* differentiated cells following treatment. Data is presented as mean±SEM of 3 independent experiment. *=*P*<0.05.

Overall, differentiated SH-SY5Y overexpressing *3K-SNCA* showed a different pattern of response to the compounds compared to the undifferentiated cells. The PIKfyve inhibitor YM201636 was able to induce a statistically significantly reduction in a-synuclein aggregates after 24 hours of treatment. To confirm that YM201636 was acting through a TFEB-mediated pathway, we investigated the levels of nuclear TFEB at different time points following neuronal differentiation. Our results confirm that YM201636 treatment was able to significantly increase nuclear TFEB levels. However, in this case we observed that the effect was more rapid than in undifferentiated cells and was established at 7 hours, with no difference recorded at 24 hours (Fig. 8, a-c). Subsequent investigation on GCase levels and activity did not detect any significant increase in GCase cytoplasmatic levels, with a non-significant trend towards increased enzyme activity observed only after 24 hours treatment (Fig. 9, a-c).

**Figure 8.**
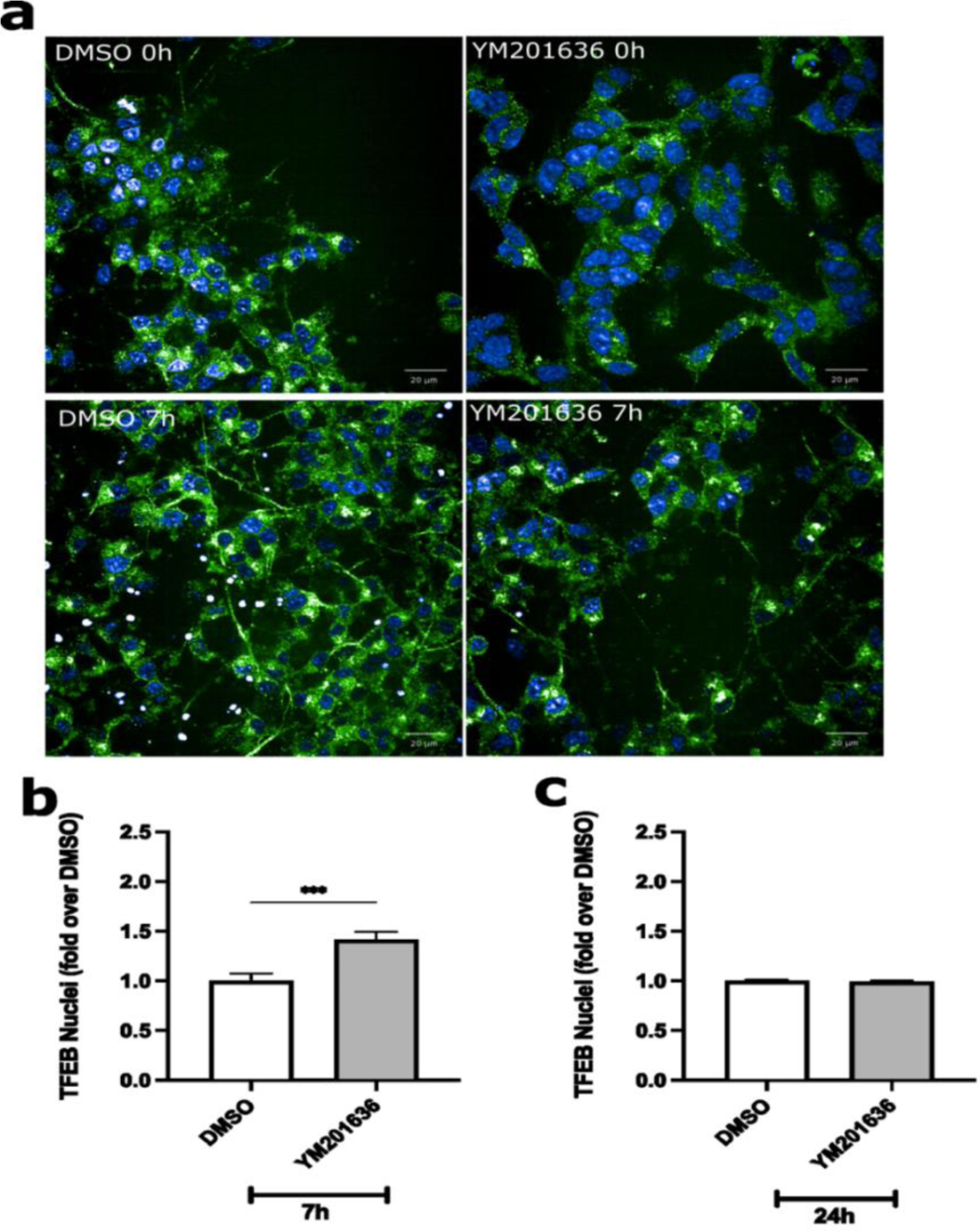
TFEB increased translocation into the nuclei following YM201636 treatment. **a)** Representative confocal images showing increased TFEB into the nuclei after 7 hours treatment. **b)** Quantification of TFEB nuclei signal intensity after 7 hours treatment **c)** Quantification of TFEB nuclei signal intensity after 24 hours treatment. Data are presented as mean±SEM of 3 independent experiment. ***=*P*<0.001.

**Figure 9.**
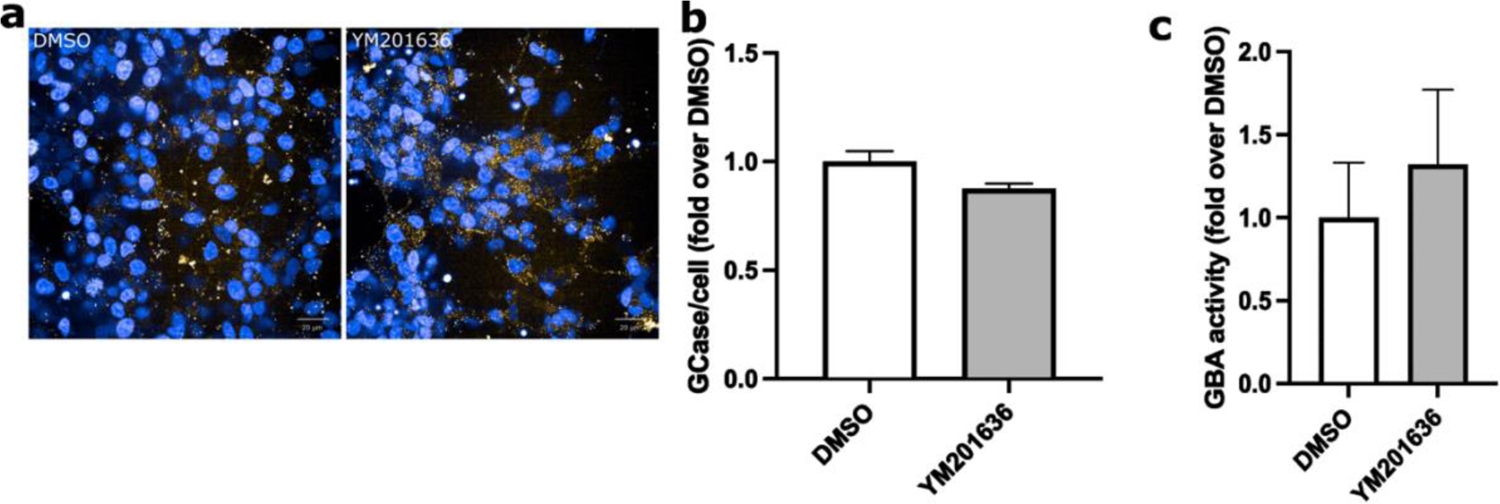
Investigation on GCase quantity and activity in differentiated cells following treatment. **a)** Representative images showing GCase levels after compounds exposure. **b)** Quantification of GCase spots after 24 hours treatment. **c)** Endpoint GCase enzyme activity quantification. Data is presented as mean±SEM of 3 independent experiment.

Finally, LAMP1 punctate quantification in differentiated 3K-SNCA cells after PIKfyve inhibition showed a statistically significant after 24 hours (Fig. 10, a-b).

**Figure 10.**
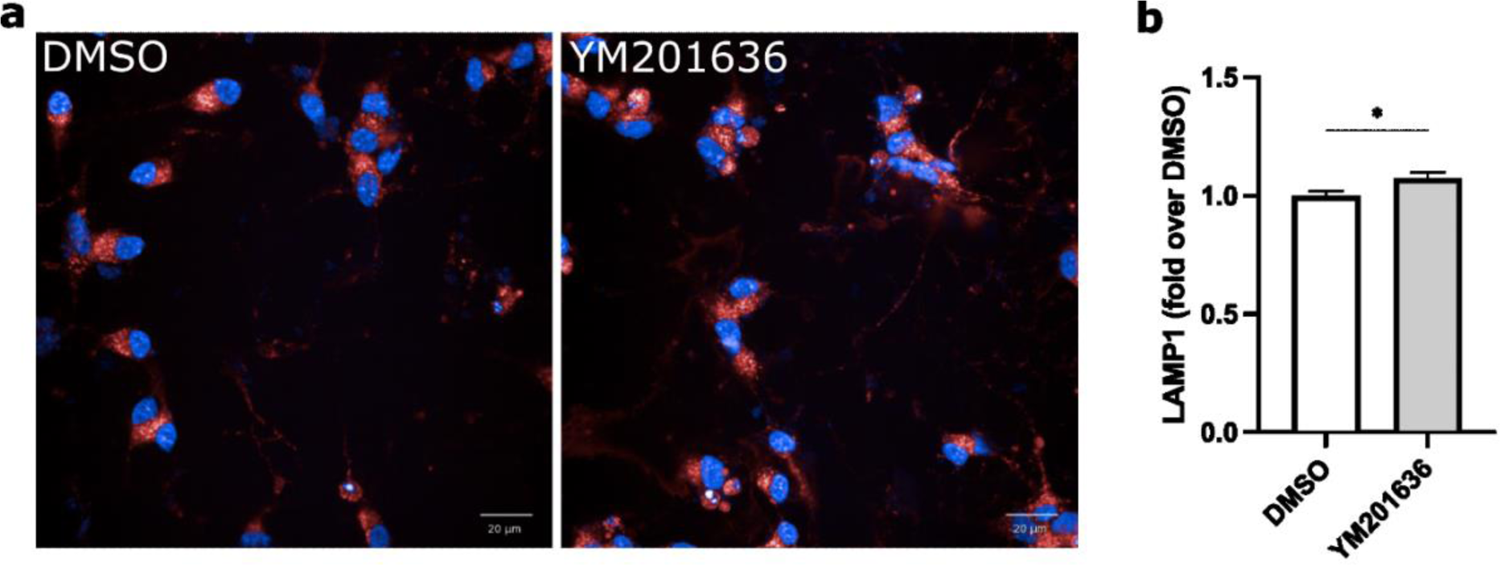
YM201636 treatment increase LAMP1 presence per cell in differentiated 3K-SNCA cells when compared to vehicle control. a) Representative images showing increased LAMP1 levels after compounds exposure. b) Quantification of LAMP1 after 24 hours treatment. Data is presented as mean SEM of 3 independent experiment. *=P<0.05.

Together, these results indicate that a-synuclein induced lysosomal dysfunction occurs in differentiated *3K-SNCA* cells. The PIKfyve inhibitor YM201636 treatment was capable of enhancing TFEB nuclear translocation in the differentiated cells model, resulting in an increase in LAMP1 levels. However, without showing any statistically significant change in GCase levels or activity was shown.

## Discussion

Lysosomal dysfunction has been implicated in the pathogenesis of PD ^67^. In post-mortem brain tissue of PD patients, there is evidence of a reduction in lysosomal enzyme activity and increased lysosomal membrane permeability ^12^. Evidence from genetic studies has confirmed the significant role of the lysosome in the pathogenesis of PD. ^11^.Variants in the *GBA1* gene are numerically the most important genetic risk factor for PD described to date ^6,68^. Additionally, mutations in the Leucine-Rich Repeat Kinase 2 (*LRRK2*) gene, which encodes a protein involved in the regulation of lysosomal trafficking, are the most common cause of familial PD, although with variable penetrance ^69,70^. Several studies have reported the relationship of lysosomal function to a-synuclein misfolding, aberrant clearance and ultimately its toxic aggregation ^22,57,59,67,71–73^.

The lysosome plays a critical role in the clearance of a-synuclein, since a-synuclein is physiologically degraded by both the ubiquitin-proteasome system and the autophagy-lysosomal pathway ^74^, mainly chaperone-mediated autophagy (CMA) and macroautophagy ^75–78^. As a result, it is anticipated that abnormality of lysosomal activity may impair the turnover of a-synuclein and facilitate an increase in a-synuclein levels. Alternatively, a-synuclein accumulation has been noted to reduce GCase activity in the absence of *GBA1* variants, leading to further impairment of a-synuclein turnover ^22,59,79^.

In this study, we have showed that overexpression of *3K-SNCA* that leading to formation of a-synuclein aggregates compromises wild-type GCase activity. Undifferentiated SH-SY5Y cells overexpressing *3K-SNCA* show a significant increase of a-synuclein aggregates and reduced GCase activity compared to the wild-type control. This effect persists after differentiation of the cells to a neuron-like phenotype. Gegg et al.^12^ showed a decrease in both GCase activity and protein levels by 70% and 87%, respectively, in human SH-SY5Y overexpressing high levels of exogenous a-synuclein. Moreover, hippocampal and cortical neurons as well as differentiated dopaminergic SH-SY5Y revealed decreased GCase activity when treated with preformed a-synuclein fibrils (PFF)^18,80^. More recent studies in a-synuclein overexpressing cell models have identified accumulation of a-synuclein at the ER, leading to ER fragmentation and aberrant accumulation and aggregation of immature GCase protein in the ER, causing an eventual functional decrease of GCase constitutional activity^24^. In conclusion, our results confirm and extend previous reports, reporting that a-synuclein aggregation and accumulation causes lysosomal dysfunction. Simultaneously, reduced GCase activity enhances pre-existing a-synuclein aggregation.

Next, we attempted to prevent the pathological a-synuclein aggregation by treating the *3K-SNCA* cells with compounds that are reported to enhance lysosomal biogenesis via TFEB modulation, in particular PIKfyve inhibitor YM201636. Our results showed that PIKfyve inhibition resulted in an upregulation of TFEB, with a reduction of a-synuclein aggregates in a lysosomal-dependent manner.

Previous *in vivo* data in rodent models suggested that overexpression of a-synuclein induced dynamic changes in TFEB, initially promoting its nuclear translocation, but sequestering TFEB into the cytoplasm when cell death occurs ^33^. This modification in TFEB subcellular localisation correlated with a progressive decline in markers of lysosome function. Interestingly, TFEB colocalises with a-synuclein and has also been observed in Lewy body-containing nigral neurons in human PD brains ^33^. Transgenic technology to induce TFEB overexpression as well as pharmacological activation of TFEB are able to potentially restore lysosomal biosynthesis and reduce the accumulation of aggregated a-synuclein by promoting its autophagic clearance ^81^.

In our model of undifferentiated cells, the increase in a-synuclein observed in undifferentiated *3K-SNCA* cells was partially reversed following a short-duration treatment with PIKfyve inhibition by YM201636. The mechanism of action of the compound was reliant on lysosomal function, as no reduction of a-synuclein aggregates was achieved when compounds were co-administrated with bafilomycin-A1. Additionally, PIKfyve inhibitor treatment in undifferentiated cells resulted in an increase in nuclear TFEB to a similar extent of the control drug ambroxol, whose signal was significantly increased in the nuclei after 24 hours. In conclusion, in our study, the observed TFEB nuclear presence, mediated by PIKfyve inhibition, was followed by an increase of lysosomal markers LAMP1 and GCase, further demonstrating the lysosomal-dependent mechanism of YM201636.

In summary, cytoplasmatic TFEB was incorporated into the nucleus, so that it executed its transcriptional activity, causing an eventual increase in lysosomal markers, supporting the central role of TFEB in protecting the cell against a-synuclein accumulation.

Although SH-SY5Y cells constitute a valuable asset to study the molecular complexity of PD, this cell line was obtained as a neuroblastoma derivative and, as a consequence, it possess physiological characteristics which differ greatly from the constitutive dopaminergic neuronal properties in differentiation fate, viability, growth performance, metabolic properties and genomic stability ^82^. One of the main limitations of SH-SY5Y is its active cellular division, which results in changes in lysosomal activity and composition, opposite to what is observed in neuronal cells where a constant G0 phase of the cell cycle is present ^83^. To overcome this limitation, *3K-SNCA* cells were differentiated into neuron-like phenotype, and this procedure did not affect the presence of a-synuclein aggregates, indicating that a-synuclein induced lysosomal dysfunction still occurs in differentiated 3K-SNCA cells. PIKfyve inhibition by YM201636 resulted in a significant reduction of a-synuclein aggregates and upregulation in TFEB nuclear incorporation. In line with the increase in TFEB nuclear levels, LAMP1 levels in the cells were also elevated, supporting the fact that PIKfyve inhibition results in TFEB translocation to the nucleus, and, eventually, in the upregulation of the TFEB targets. Nevertheless, GCase protein levels and activity did not show significant difference when compared to control. This lack of effect in GCase could be attributed to enhanced aggregation of a-synuclein at the ER, causing ER fragmentation, compromising folding capacity and aggregation of lysosomal hydrolases in the ER^24^ and, hence, causing accumulation of immature forms of GCase in the ER and reduction of protein levels and functionality.

PIKFyve inhibition has been suggested as a therapeutic target for several neurodegenerative diseases. Improvement in the survival rate of motor neurons derived from iPSC with ALS caused by a repeat expansion in C9ORF72 was found after treatment with YM201636 ^84^ and the pharmacological inhibition with YM201636 decreased the lysosomal delivery of tau aggregates, halting a key step in the progression of tauopathies ^85^. In our study, PIKfive inhibitor was capable of reducing a-synuclein aggregates and increasing TFEB nuclear translocation both in undifferentiated and differentiated cells. In undifferentiated cells, YM201636 effect was, at least to some extent, dependent on the lysosomal function. However, this was not translated into elevated GCase levels or activity. One possible explanation could be that, although YM201636 stimulated TFEB, this stimulation did not fully compensate for the GCase reduction induced by a-synuclein accumulation.

In differentiated cells, YM201636 was capable of enhancing TFEB nuclear translocation. This result may be attributed to PIKfyve positively regulation of mTORC1 phosphorylation on TFEB, confining it outside of the nucleus ^86^. Following PIKfyve inhibition, phosphatase 2 A dephosphorylates TFEB on Ser211 enabled its translocation into the nucleus, thus activating the CLEAR network ^42^. Additionally, it has been proposed that PIKfyve could impact on TFEB independently of mTORC1 via its interaction the serine/threonine kinase Akt (protein kinase B), which modulates TFEB by phosphorylating it at Ser467 and represses TFEB nuclear translocation ^87^. YM201636 has been proposed to inhibit Akt phosphorylation in adipocytes ^88^, and PIKFfYve inhibitors cytotoxicity in fibroblast is evident only when the PIKfyve inhibitor is combined with AKT signalling pathway suppression ^89^. However, to determine whether potentially YM201636 could modulate TFEB via alternative mechanism has not been entirely established and requires further research.

Our results suggest that YM201636 modulates phosphorylation on TFEB, ultimately resulting in reduction of a-synuclein aggregates in *3K-SNCA* model, presumably by promoting the lysosomal-autophagy pathway.

## Conclusion

This work demonstrates the potential role that the TFEB pathway presents as a therapeutic target in the treatment of neurodegenerative diseases characterised by the accumulation of misfolded proteins. Our results identify PIKfyve signalling as a target to increase TFEB-mediated lysosomal biogenesis, resulting in a reduction of a-synuclein aggregates in *3K-SNCA* SH-SY5Y and neuron-like cells.

## Data availability

Data and methods that support the findings of this study will become openly available at https://zenodo.org and https://www.protocols.io repositories respectively.

## Financial disclosures of all authors (for the preceding 12 months)

The authors have no financial disclosures to declare for the preceding 12 months.

## Acknowledgments

This research was funded by Aligning Science Across Parkinson’s [Grant number: ASAP-000420] through the Michael J. Fox Foundation for Parkinson’s Research (MJFF). For the purpose of open access, the author has applied a CC BY 4.0 public copyright license to all Author Accepted Manuscripts arising from this submission.

## Abbreviations

AD: Alzheimer disease

ALS: Amyotrophic lateral sclerosis

a-synuclein: Alpha-synuclein

BDNF: Brain-derived neurotrophic factor

CLEAR: Coordinated Lysosomal Expression and Regulation gene network

CMA: Chaperone-mediated autophagy

CMT4J: Charcot-Marie-Tooth syndrome

ER: Endoplasmatic reticulum

FBS: Fetal bovine serum

GCase: glucocerebrosidase

GlcCer: Glucosylceramide

GlcSph: Glucosylsphingosine

hiPSC: Human induced pluripotent stem cells

HLH: helix-loop-helix

LAMP1: Lysosomal Associated Membrane Protein 1

LRRK2: Leucine-Rich Repeat Kinase 2 gene

mTORC1: Mechanistic target of rapamycin complex 1

NGS: Normal goat serum

PBS: Phosphate-buffered saline buffer

PD: Parkinson disease

PFA: Para-formaldehyde

PFF: Preformed a-synuclein fibrils

pS129: phosphorylated serine residue 129

RA: Retinoic Acid

Ser142, Ser211: Serine residues 142 and 211

SNpc: Substantia nigra pars compacta

TFEB: Transcription Factor EB TFEB

TRPML1: Mucolipin TRP channel

## Supplementary figures

**Figure S1.**
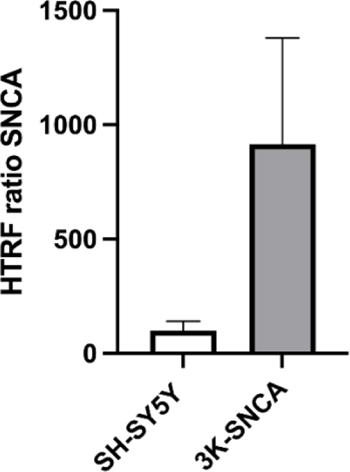
HTRF-FRET quantification of α-synuclein aggregates in parental SH-SY5Y and 3K-SNCA overexpressing cells.

**Figure S2.**
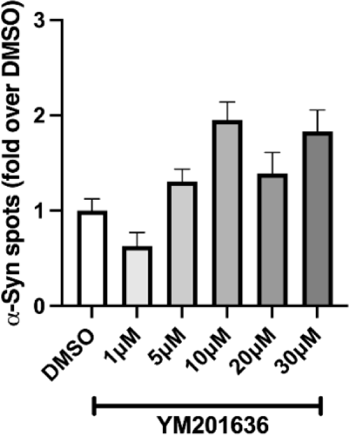
Serial dilution pilot experiment to identify optimal concentration of the various compounds able to reduce synuclein level. Data is presented as mean ± SEM.

**Figure S3.**
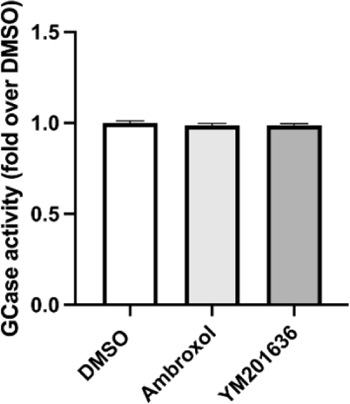
GCase activity quantification following 24 hours treatment with different compounds. Data is presented as mean ± SEM.

**Figure S4.**
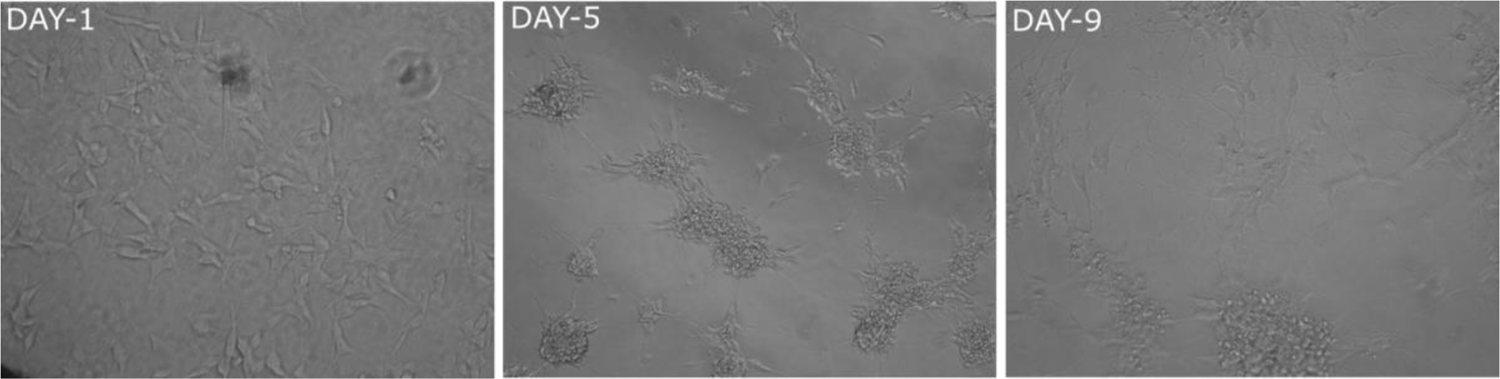
Stages of SH-SY5Y differentiation into neuron-like cells. Axons can be observed emerging, along with classic clumping of cell bodies.

