## Supplementary figures and images for "Reduction of a-synuclein aggregates by PIKfyve inhibition via TFEB-mediated lysosomal biogenesis in a Parkinson disease model"

### Supplementary figure 1

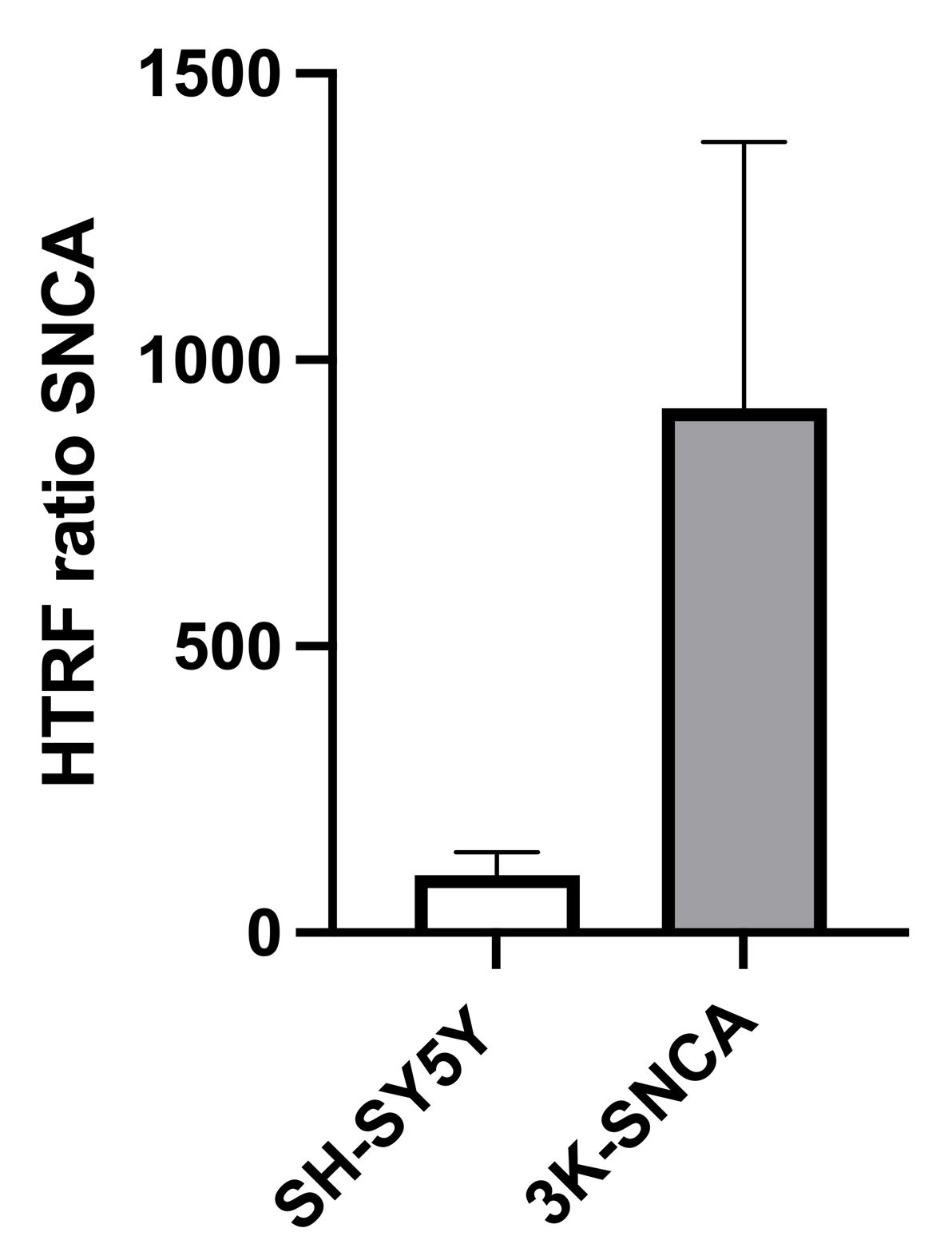

### Supplementary figure 2

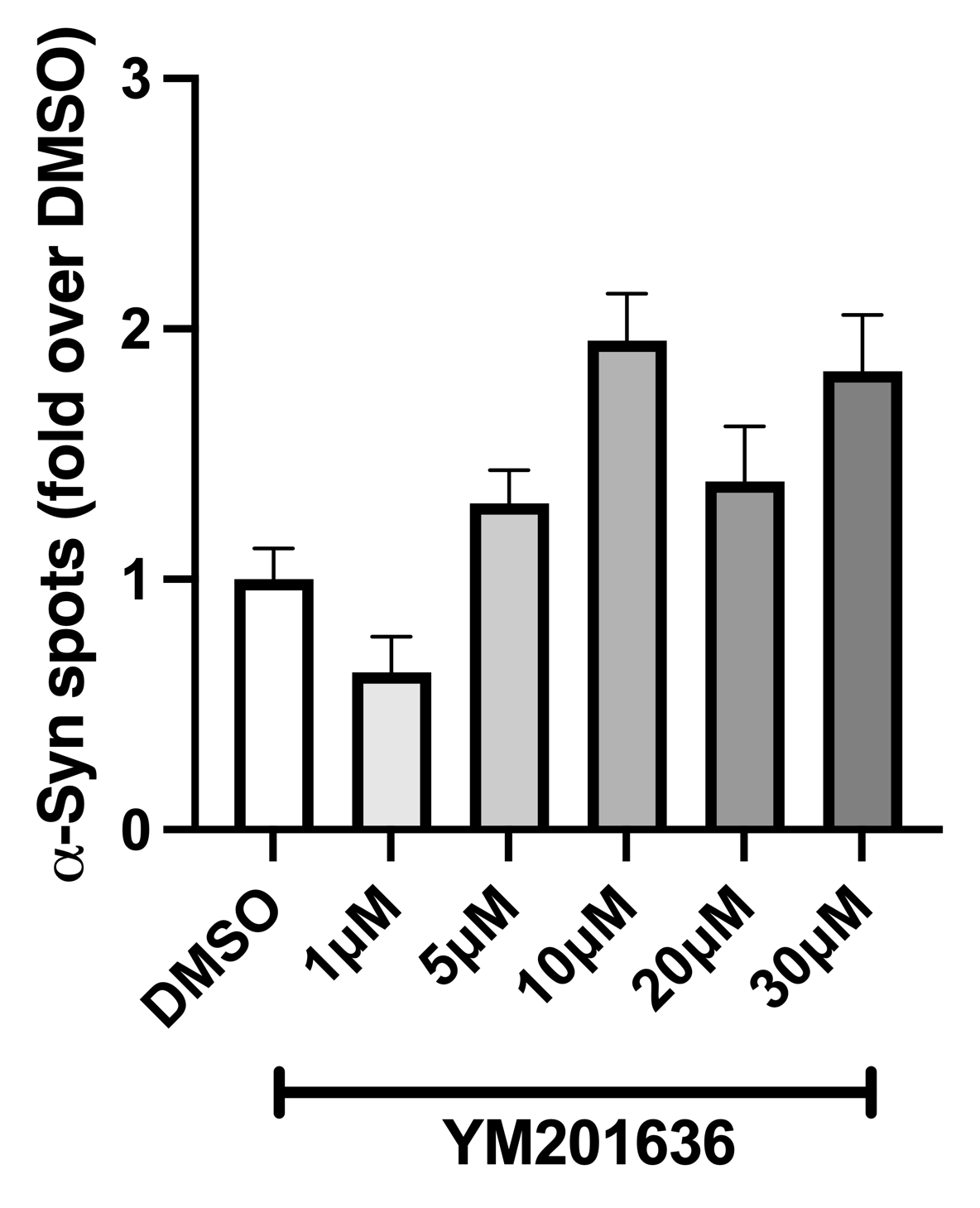

### Supplementary figure 3

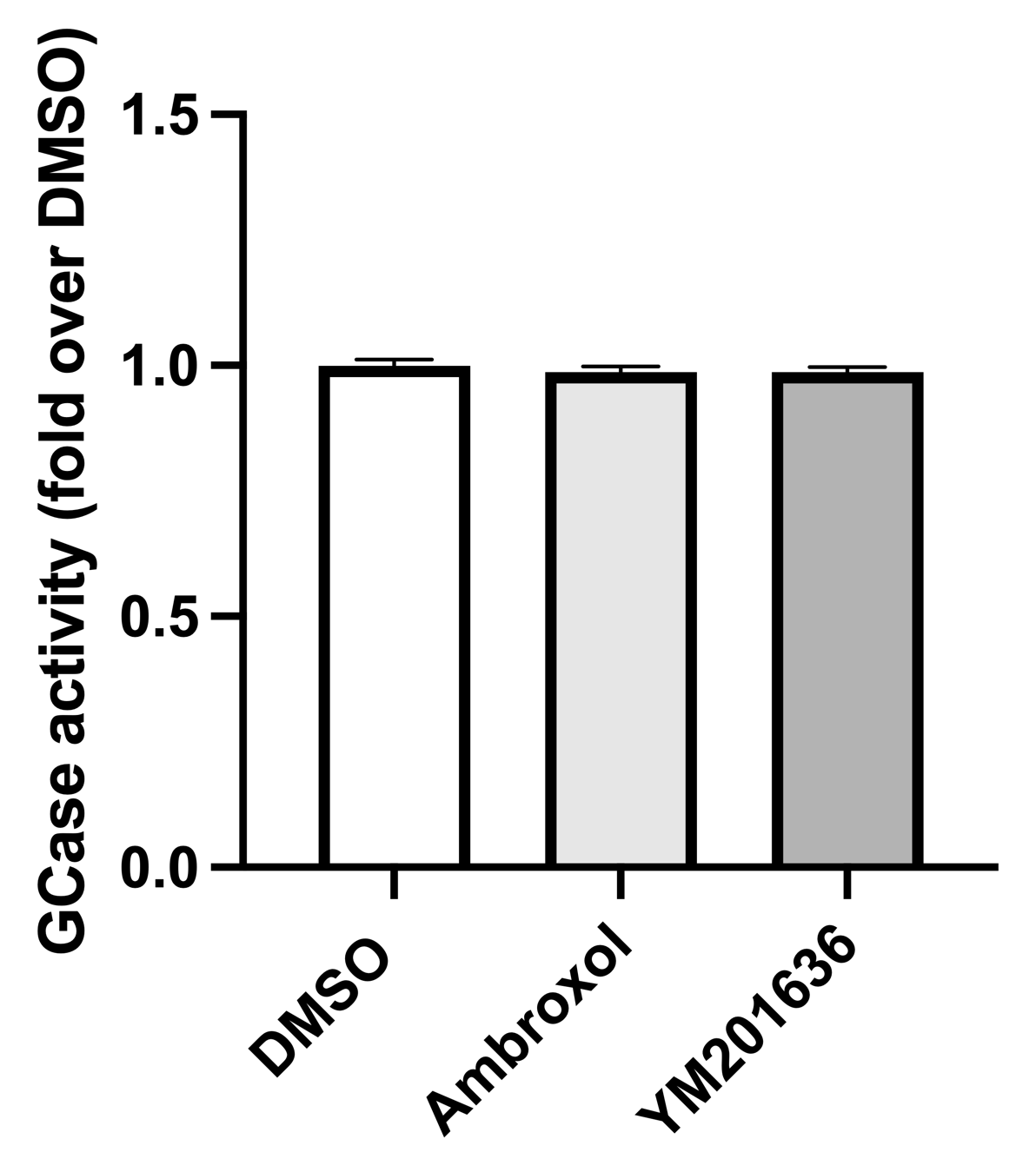

### Supplementary figure 4

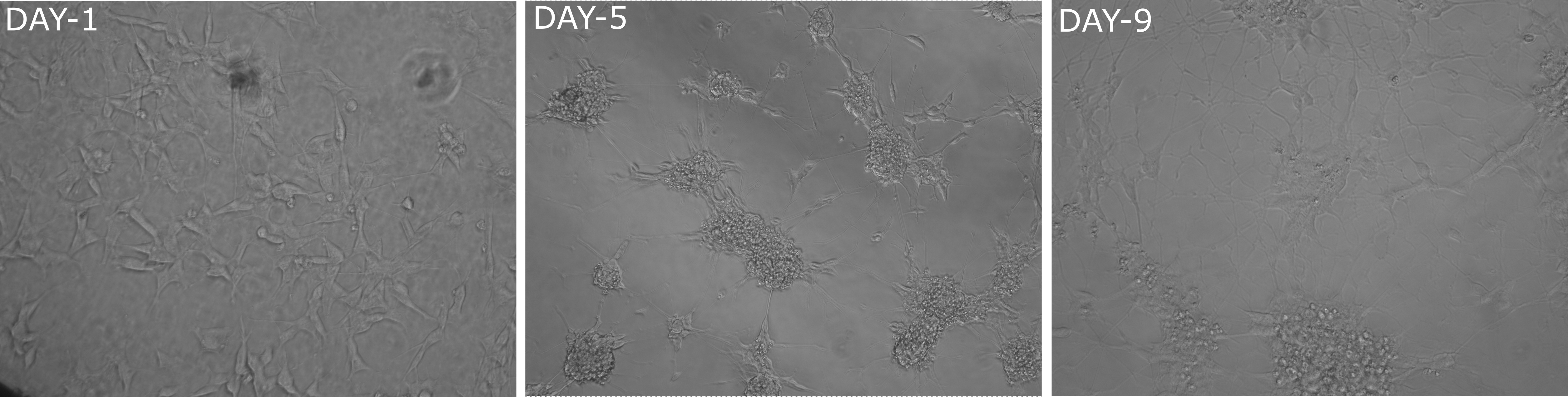
